# Plscr1 inhibits murine norovirus entry

**DOI:** 10.64898/2026.09.17.752154

**Authors:** Nina S. Baggett, Linley R. Pierce, Robert C. Orchard

## Abstract

Noroviruses are a leading cause of gastroenteritis worldwide, yet host factors that restrict norovirus replication are not well understood. By mining both CRISPR activation and CRISPR knockout genome-wide screens, we identified the interferon-stimulated gene Plscr1 as a restriction factor for murine norovirus (MNV). Plscr1 inhibits the fusion of enveloped viruses with endocytic membranes, making its antiviral activity against the nonenveloped MNV surprising. Here, we demonstrate that Plscr1 is both necessary and sufficient to restrict MNV infection in vitro and contributes to the control of colonic infection in mice. Mechanistically, we determined that Plscr1 inhibits MNV entry at a post-attachment step. A single amino acid substitution in the minor capsid protein VP2 confers resistance to Plscr1. Because VP2 delivers viral RNA by puncturing endocytic membranes during calicivirus entry, these findings implicate a late entry step targeted by Plscr1. Our results expand the antiviral activity of PLSCR1 to a nonenveloped virus and identify an unexpected point of convergence between enveloped-and non-enveloped-virus entry pathways.

## Introduction

Noroviruses are non-enveloped, positive-sense, single-stranded RNA viruses that are the leading cause of viral gastroenteritis^1,2^. Noroviruses pose a substantial global health and economic burden, with an estimated annual societal cost of $10.6 billion in the United States alone^3^. Our knowledge of human norovirus (HNoV) is limited due to the historical challenges in cultivating the virus in vitro and lack of a small animal model. The barrier to cultivating HNoV in vitro has been overcome with the discovery that nontransformed human intestinal enteroids (HIEs) can support HNoV replication^4^. Several challenges persist in establishing robust HNoV growth in vitro that limit the ability to interrogate HNoV biology^5^. However, recent advances, including the generation of infectious molecular clones and a protocol for creating stocks of some HNoV strains are promising developments^6,7^. Murine norovirus (MNV) is a natural pathogen of mice that replicates robustly in cell lines, providing a platform to investigate norovirus biology both in vitro and in vivo in a natural setting. However, MNV is an imperfect human norovirus model, as mice lack the capacity to vomit, infection rarely leads to diarrhea, and MNV encodes an additional open reading frame^8^. Despite these differences, both HNoV and MNV share many significant features including fecal-oral transmission, infection of the intestines with fecal shedding, primary genome organization, and molecular mechanisms of RNA expression and replication^8^. These shared aspects have empowered the MNV model to uncover fundamental norovirus biology.

MNV strains share high genetic similarities but exhibit phenotypic diversity in mice, driven largely by differential sensitivity to interferon-mediated restriction^9,10^. For example, type I IFN plays a major role in restraining MNV strain CW3 (MNV^CW3^) infection, as infection of IFN-α/βR^−/−^ mice with MNV^CW3^ is lethal^11^. Additionally, type I and type III IFN signaling prevent MNV^CW3^ infection of tuft cells^12,13^. HNoV strains also show strain-dependent differences in sensitivity to IFN^14^. These data highlight the importance of IFN in shaping norovirus tropism and pathogenesis. However, the downstream interferon-stimulated genes (ISGs) that mediate norovirus restriction are not known and their identification would provide insight into norovirus biology and innate immunity.

Multiple genome-wide CRISPR/Cas9 screens have been conducted to identify proviral and antiviral genes involved in MNV infection^15-19^. By mining previously published genetic screening data to identify broadly acting anti-norovirus genes, we identified phospholipid scramblase 1 (PLSCR1) as an MNV restriction factor. PLSCR1 is an ISG induced by type I, II, and III interferons^20,21^. PLSCR1 restricts many viruses including the entry of HIV and SARS-CoV-2^21-23^. The prevailing model is that PLSCR1 blocks the fusion between viral envelopes and endomembranes^21-23^. However, it remains unclear whether PLSCR1 can restrict non-enveloped viruses such as MNV. Here, we demonstrate that Plscr1 is both necessary and sufficient to restrict MNV replication in vitro. Mice deficient in *Plscr1* have a modest MNV replication phenotype in the colon. PLSCR1 inhibits MNV entry and this restriction can be overcome with variation in the minor capsid protein VP2. Taken together, our data point to a critical role for PLSCR1 in restricting MNV replication and expands PLSCR1’s antiviral activity to include non-enveloped viruses.

## Results

### PLSCR1 is sufficient to restrict MNV replication

We mined three previously conducted genome-wide CRISPR screens to systematically identify candidate antiviral genes^16,17,19^. The screens are all pooled screens using cellular survival after MNV infection as a selective pressure, but differ in cell type and perturbation strategy. First, we included a high density CRISPR knockout screen performed in BV2 cells, a murine microglial cell line that is naturally susceptible to MNV infection. This loss-of-function screen identified candidate antiviral genes whose knockout increased sensitivity to MNV infection, a negative selection strategy^19^. The second screen used a genome-wide CRISPR activation (CRISPRa) approach in BV2 cells in which the overexpression of putative antiviral genes would provide a growth advantage to cells when challenged with MNV^16^. The final screen we included in our analysis was also a CRISPRa screen but conducted in HeLa cells expressing the MNV receptor, CD300lf^17^. Our analysis from these three screens consistently identified two genes with antiviral activity towards MNV: IFITM2 and PLSCR1 (**Figure 1A**). Both IFITM2 and PLSCR1 are interferon stimulated genes with known antiviral roles targeting the entry of enveloped viruses. IFITM2 is part of the interferon-inducible transmembrane (IFITM) protein family which blocks the entry of viruses from the endomembrane compartment^24^. While classically investigated in the context of enveloped virus infection, IFITM3 inhibits reovirus, a non-enveloped virus^25^. Phospholipid scramblase 1 (PLSCR1) is an ISG induced by all three types of IFN and blocks the fusion of HIV and SARS-CoV-2 with endocytic membranes^21-23^. Besides blocking entry, other enveloped viruses are inhibited by PLSCR1 through distinct mechanisms^20,26,27^. To date, PLSCR1 has not been shown to inhibit any non-enveloped virus. Therefore, we have elected to focus on how PLSCR1 restricts MNV replication, potentially revealing new aspects of norovirus biology and unappreciated mechanisms of PLSCR1 restriction.

**Figure 1:**
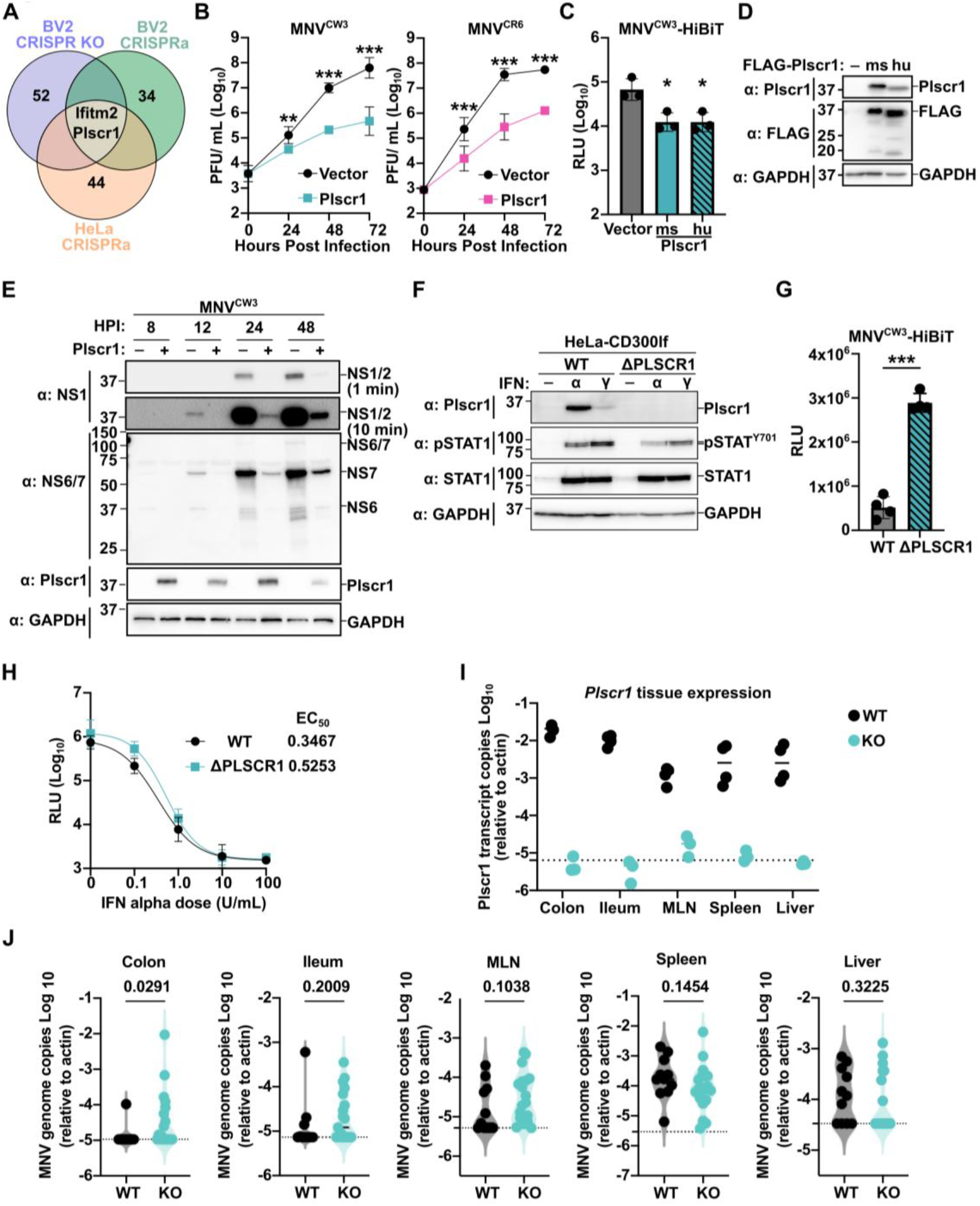
Plscr1 is required for optimal control of MNV infection. (**A**) Venn diagram depicting conserved hits across three distinct screens. (**B**) HeLa-CD300lf cells expressing either an empty vector or Plscr1 were infected with MNV^CW3^ (left) or MNV^CR6^ (right) at a multiplicity of infection (MOI) of 0.05. Viral production was enumerated using plaque assays (PFU; plaque forming units). (**C**) HeLa-CD300lf cells expressing mouse or human Plscr1 were challenged with MNV^CW3^-HiBiT at a MOI of 0.5 for 48 hours. Infection was quantified using Nano-Glo HiBiT Lytic Detection System (RLU; relative luminescent units). (**D**) Representative western blot of HeLa-CD300lf cells expressing FLAG-tagged mouse (ms) or human (hu) Plscr1 probed with indicated antibodies. (**E**) Representative western blot of HeLa-CD300lf cells expressing either vector control or Plscr1 infected with an MOI of 5.0 of MNV^CW3^ and lysed at indicated hours post-infection (hpi). (**F**) Representative western blot of HeLa-CD300lf or HeLaΔPlscr1-CD300lf cells treated with no interferon, 1000 units/mL interferon α, or 0.1 μg/mL interferon γ probed with indicated antibodies. (**G**) HeLa-CD300lf or HeLaΔPlscr1-CD300lf cells were challenged with an MOI of 0.5 of MNV^CW3^-HiBiT. After 48 hours, infection was quantified using Nano-Glo HiBiT Lytic Detection System. (**H**) HeLa-CD300lf or HeLaΔPlscr1-CD300lf cells were treated with indicated doses of interferon α (U/ml) then challenged with MNV^CW3^-HiBiT at a MOI of 0.5 for 48 hours. Infection was quantified using Nano-Glo HiBiT Lytic Detection System. Effective concentration 50 (EC_50_) was calculated for interferon α for indicated cell types. (**I**) *Plscr1* expression from *Plscr1*^+/+^ (WT) or *Plscr1*^-/-^ (KO) littermates in indicated tissues. Dashed line indicates limit of detection.(**J**) *Plscr1*^+/+^ (WT) or *Plscr1*^-/-^ (KO) littermates were infected with 1 × 10^6^ PFU of MNV^CW3^ and euthanized 3 days post-infection. Colon, Ileum, Mesenteric Lymph Nodes (MLN), spleen and liver were analyzed via qPCR for MNV genome copies and normalized to actin. Individual data points are shown along with a violin plot for each tissue. P-values from Mann-Whitney test are shown. Dashed line indicates limit of detection. All data are mean ± standard deviation from three independent experiments and analyzed by one-way ANOVA with Tukey’s multiple comparison test (**B and C**), T-test (**G)**, or Mann-Whitney (**J**). *P < 0.05, **P < 0.01, ***P < 0.001, ****P < 0.0001.

To test if PLSCR1 is sufficient to inhibit MNV replication, we assessed the ability of MNV to replicate when PLSCR1 is overexpressed in HeLa-CD300lf cells (**Figure 1B**). We chose HeLa-CD300lf cells because they were one of the cell lines used in the initial screen and because they are more genetically tractable than mouse macrophage-like cells. While two strains of MNV (MNV^CW3^ and MNV^CR6^) replicated robustly over 72 hours in vector control cells, expression of mouse Plscr1 significantly attenuated viral growth (**Figure 1B**). Although mouse and human PLSCR1 proteins share only 75% sequence identity, overexpression of either mouse or human PLSCR1 inhibits MNV replication in HeLa-CD300lf, as measured using a recently developed split-luciferase reporter of MNV (MNV-HiBiT)^28^ (**Figure 1C and 1D**). Consistent with the decrease in viral production, Plscr1 expressing cells had scant levels of viral protein production after MNV challenge compared to vector control cells (**Figure 1E**). Taken together, these data indicate that PLSCR1 is sufficient to inhibit MNV replication.

### Plscr1 is necessary for restricting MNV in Vitro

To determine if PLSCR1 is necessary for optimal restriction MNV, we generated HeLaΔPLSCR1 cells that express CD300lf (HeLaΔPLSCR1-CD300lf). We confirmed PLSCR1 was knocked out by stimulating cells with IFN-α or IFN-γ and detecting PLSCR1 expression via western blot on lysates from HeLa-CD300lf cells and not HeLaΔPLSCR1-CD300lf (**Figure 1F**). Consistent with our screening data, when challenged with MNV-HiBiT we observed an increase in viral replication in HeLaΔPLSCR1-CD300lf cells compared to HeLa-CD300lf (**Figure 1G**). Prior work suggests that PLSCR1 may amplify the type I IFN response while other studies indicated that PLSCR1 is a direct-acting antiviral^20-23,26,27^. To test if the antiviral role of PLSCR1 is potentiation of IFN signaling, we compared MNV sensitivity to type I IFN in the presence or absence of endogenous PLSCR1. IFN-α treatment significantly reduced MNV replication in a dose-dependent manner regardless of PLSCR1 expression (**Figure 1H**). We observed no significant differences between the EC50 values in these cell lines, indicating that PLSCR1 deficiency does not alter the type I IFN response to restrict MNV (**Figure 1H**). Thus, our data are consistent with PLSCR1 functioning as a direct antiviral molecule that is necessary for optimal control of MNV.

### Plscr1 deficient mice have a modest increase in MNV colonic replication

Interferon signaling shapes infection dynamics of MNV in vivo^11-13,29-32^. However, the specific ISG effectors that mediate interferon-dependent restriction of MNV infection in vivo remain poorly defined. We obtained *Plscr1*^*-/-*^ mice that have been previously described^33^ and first assessed the levels of Plscr1 transcripts in tissues relevant to MNV infection from Plscr1 sufficient and deficient animals. Plscr1 transcript levels were detectable in all tissues assayed from wild type mice with the highest expression in the colon and ileum, while knockout mice had no detectable transcripts (**Figure 1I**). We orally inoculated *Plscr1* sufficient and deficient mice with MNV^CW3^ and harvested tissues 3 days post infection (**Figure 1J)**. Consistent with prior reports, we detected MNV^CW3^ genomes in the spleen, liver, and mesenteric lymph nodes (MLNs) of wild-type mice, but detected few genomes in the ileum and virtually none in the colon (**Figure 1J**). Compared to littermate controls, Plscr1 deficient mice had similar infection dynamics in the spleen, liver, and MLNs (**Figure 1J**). However, there was a significant, albeit modest, increase in MNV^CW3^ in the colon of Plscr1 deficient mice compared to wild-type littermate controls (**Figure 1J**). There was a similar trend in the ileum that was not statistically significant. These data indicate that *Plscr1* modestly suppresses MNV^CW3^ replication in the colon, suggesting that additional ISG effectors restrict MNV replication and pathogenesis in vivo.

### A forward genetic screen identifies a VP2 substitution confers Plscr1 resistance

To determine how Plscr1 restricts MNV, we performed a forward genetic screen in which MNV^CR6^ adapts to growth on HeLa-CD300lf expressing Plscr1 or control cells (**Figure 2A**). After 11 passages, the virus passaged on HeLa-CD300lf cells expressing Plscr1 (Plscr1 p11) displayed a growth advantage over virus passaged on vector control HeLa-CD300lf cells (Vector p11**; Figure 2B**). We then identified genetic changes associated with the phenotype through amplicon sequencing of PCR products (**Figure 2C; Supplementary Table 1**). To identify potential driver mutations, we compared the mutational landscape of Vector p11 with Plscr1 p11 virus (**Figure 2D**). This comparison identified three genetic changes that were present in greater than 30% of Plscr1 p11 (**Figure 2D**). Of these alterations, one was a same-sense mutation while the other two were missense mutations (NS2^F304L^ and VP2^Q32R^; **Figure 2E)**. To determine which of the putative amino acid mutations conferred Plscr1 resistance, we introduced them into our molecular clone of MNV^CR6^ and generated infectious viruses harboring individual substitutions (MNV^CR6^ NS2^F304L^ and MNV^CR6^ VP2^Q32R^). Only MNV^CR6^ VP2^Q32R^ grew unimpaired in Plscr1-expressing cells, validating it as a bona fide Plscr1 escape mutation (**Figure 2F**).

**Figure 2:**
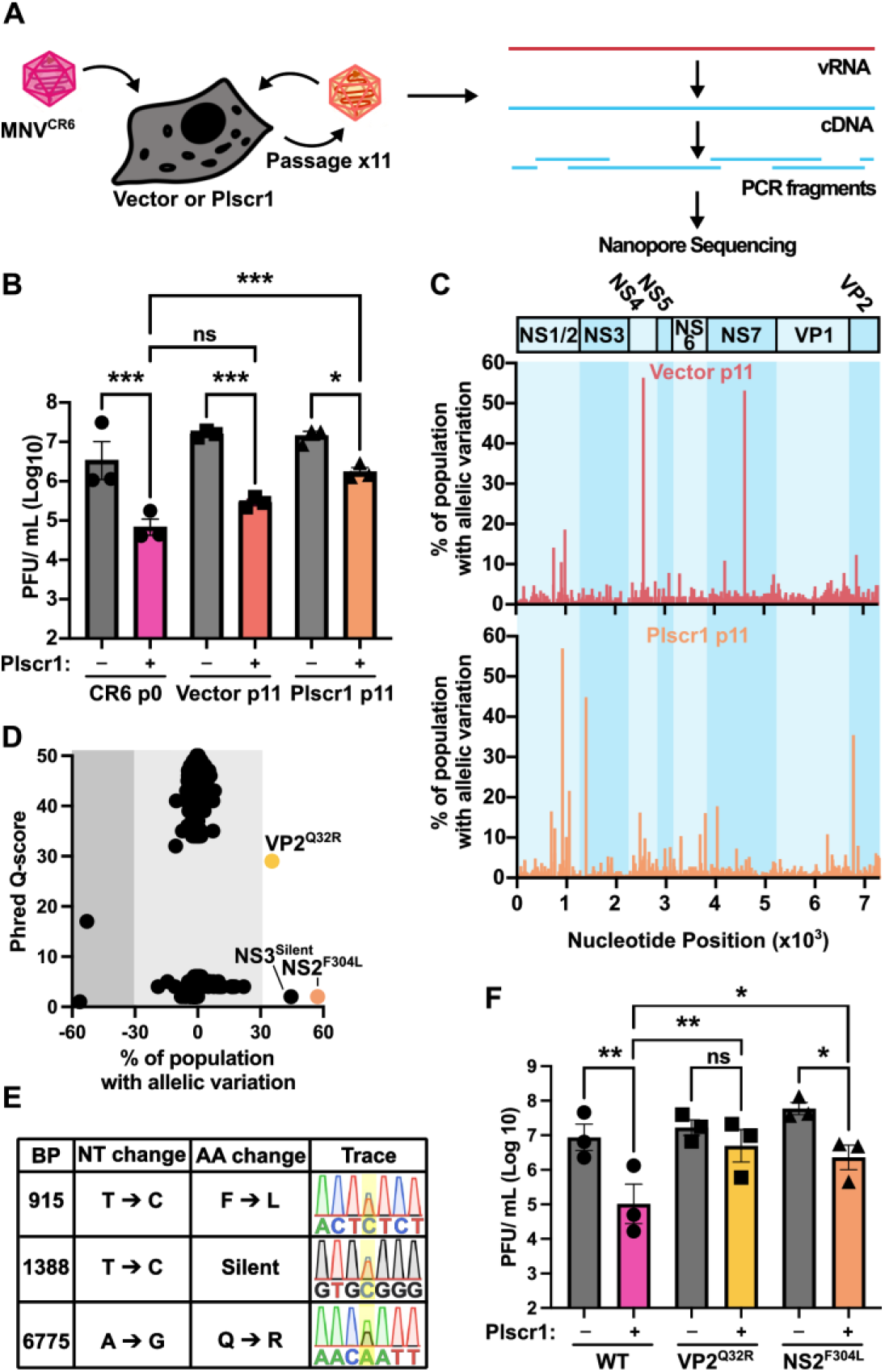
Variation within VP2 confers resistance to Plscr1 restriction. (**A**) Cartoon schematic of experimental setup to identify mutations conferring Plscr1 resistance. (**B**) HeLa-CD300lf cells expressing either an empty vector or Plscr1 were infected with parental stock of MNV^CR6^ (p0) or MNV^CR6^ p11 harvested from vector or Plscr1 expressing cells at a multiplicity of infection (MOI) of 0.05. Viral production was enumerated using plaque assays (PFU; plaque forming units) 48 hours post-infection. (**C**) Summary of sequencing coverage depicting allelic variation (y-axis) across the MNV genome (x-axis) for virus passaged 11 times on vector expressing cells (Vector p11; red coloration) or passaged 11 times on Plscr1 expressing cells (Plscr1 p11; orange coloration). Viral genome organization is shown on the top in alternating shades of blue, which is carried through the graphs. (**D**) Graph depicting allelic variation between viral populations passaged on Plscr1 compared to Vector (x-axis) with positive values being enriched in the Plscr1 p11 population. The Phred quality score for each base depicting confidence of nanopore sequence base calling for each site on the y-axis. Variants exceeding 30% of the population are labeled. (**E**) Table of the indicated variants indicating base pair (BP) position, nucleotide (NT) change, amino acid (AA) change, and the trace indicating relative variation. (**F**) HeLa-CD300lf cells expressing either an empty vector or Plscr1 were infected with indicated MNV strains derived from molecular clones at an MOI of 0.05. 48 hours post-infection, viral production was enumerated via plaque assay. All data are mean ± standard error of the mean (SEM) from 3-4 independent experiments and analyzed by one-way ANOVA with Tukey’s multiple comparison test. *P < 0.05, **P < 0.01, ***P < 0.001.

The norovirus capsid is composed of two capsid proteins: VP1 and VP2. The capsid comprises 90 dimers of VP1 (180 copies) and an unknown number of VP2 copies as it is not resolved on cryo-electron microscopy maps^34-36^. While relatively little is known about MNV VP2, the related feline calicivirus (FCV) has provided critical insight into VP2 that is likely applicable to other caliciviruses such as MNV. After receptor engagement, the FCV capsid undergoes structural rearrangements and exposes a dodecamer of VP2^37^. The VP2 complex forms a portal-like structure in which RNA is hypothesized to traverse through to enter the cytoplasm^37,38^. HNoV VP2 binds VPg (viral protein genome-linked) via the carboxy-terminal region of VP2 to facilitate assembly and presumably genome packaging^39^. Thus, the impact of Q32R on MNV VP2 portal assembly and endomembrane insertion or RNA packaging remains unclear.

### Plscr1 requires membrane-interacting regions to restrict MNV entry post attachment

VP2 has roles both in viral entry and assembly, so we conducted an entry bypass experiment to narrow down which step is inhibited in Plscr1 expressing cells. Equivalent viral titers were recovered after transfecting MNV^CW3^ viral RNA into HeLa cells expressing Plscr1 as control HeLa cells (**Figure 3A**). The rescue of viral titers in this experiment is strong evidence that Plscr1 inhibits MNV entry. We next assessed the ability of MNV to bind cells expressing Plscr1 in the presence or absence of CD300lf. Consistent with previous data, cells expressing CD300lf had a significant increase in MNV binding compared to non-CD300lf expressing cells (**Figure 3B**). Plscr1 expression had no impact on MNV binding regardless of CD300lf expression (**Figure 3B**). These data indicate that Plscr1 inhibits a post-binding entry step, consistent with data from enveloped viruses^21-23^.

**Figure 3:**
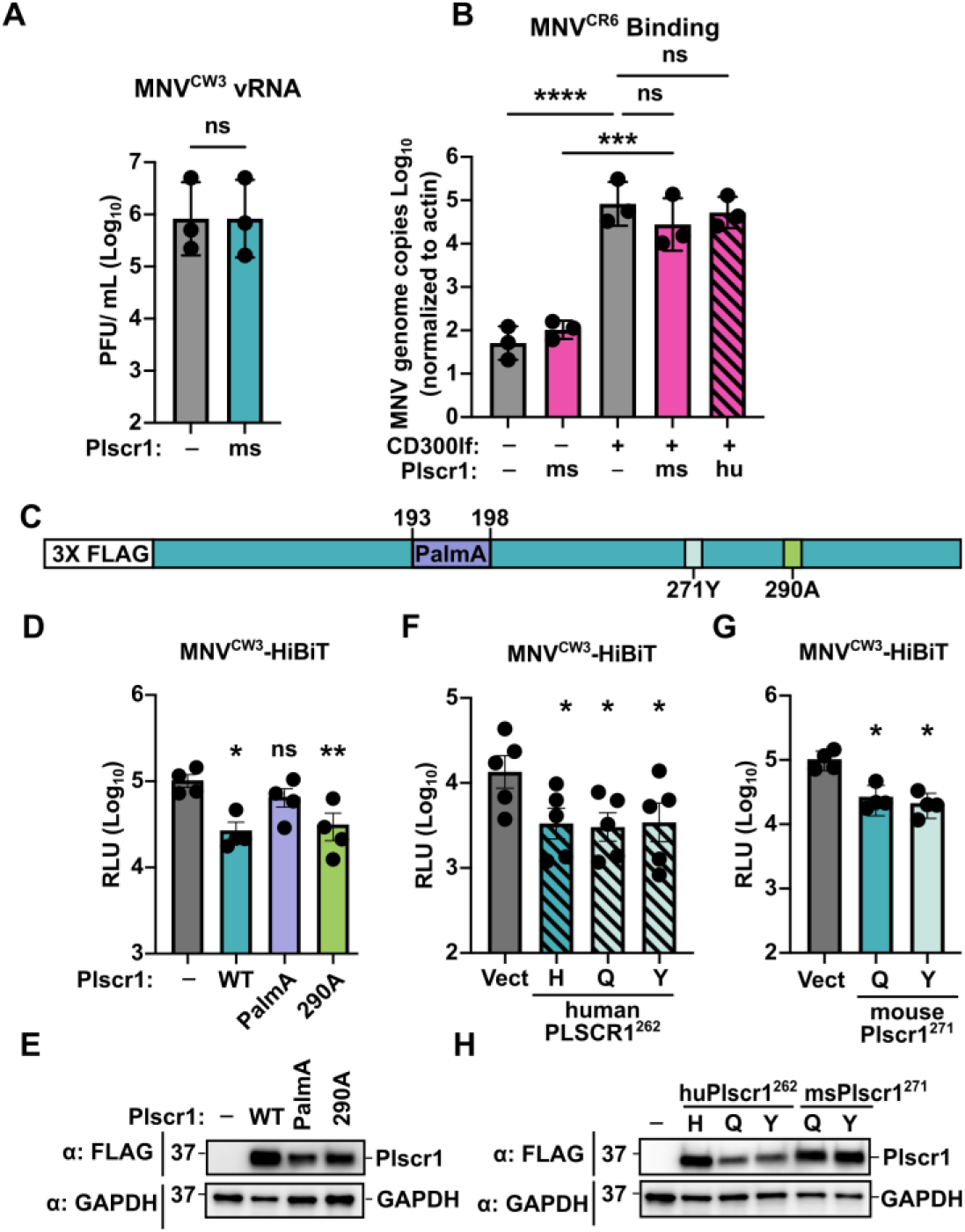
Plscr1 restricts MNV entry after viral attachment. (**A**) HeLa cells expressing empty vector or Plscr1 were transfected with 4 μg MNV^CW3^ viral RNA. 24 hours post-transfection viral titers were enumerated by plaque assay. (**B**) Indicated cell lines were assayed for MNV^CR6^ binding via qPCR. Values were normalized to account for differences in actin transcripts as detected by qPCR. (**C**) Schematic of mouse Plscr1 protein with critical regions highlighted including 3xFLAG epitope, Palmitoylation motif, equivalent polymorphism site at 271Y in mice, and previously reported scramblase mutation F290A.(**D**) HeLa-CD300lf cells expressing either an empty vector or indicated Plscr1 mutants were infected with MNV^CW3^-HiBiT at an MOI of 5. Infection was quantified 48 hours post-infection using Nano-Glo HiBiT Lytic Detection System. (**E**) Representative western blot of indicated cell lines used in (**D**). (**F-G**) HeLa-CD300lf cells expressing either an empty vector, human PLSCR1 (**F**) or mouse Plscr1 (**G**) with indicated polymorphisms at position 262 in humans or the equivalent position 271 in mice were infected with MNV^CW3^-HiBiT at an MOI of 0.5. 48 hours post-infection was quantified using Nano-Glo HiBiT Lytic Detection System. (**H**) Representative western blot of indicated cell lines used in (**F and G**). All data are mean ± standard deviation from three independent experiments and analyzed by one-way ANOVA with Tukey’s multiple comparison test. *P < 0.05, **P < 0.01, ***P < 0.001, ****P < 0.0001.

Many distinct functions have been attributed to PLSCR1 including lipid scrambling, nuclear translocation, and membrane association via palmitoylation^21,40-42^. We next assessed whether any of these previously attributed functions contribute to restriction of MNV using historically identified mutations (**Figure 3C**). Changing the amino acids reported to be critical for nuclear location ^267^<u>K</u>IS<u>KQ</u>^271^ to ^267^<u>A</u>IS<u>AA</u>^271^ diminished Plscr1 protein expression and was excluded from further analysis^43^. While originally identified as a phospholipid scramblase, Plscr1 lacks the characteristic membrane-spanning architecture of well-characterized phospholipid scramblases and thus there is debate over whether Plscr1 possesses lipid scrambling activity^21,44,45^. Plscr1^F290A^, a mutant previously implicated as a loss of scramblase function^40^, retains its ability to block MNV replication in HeLa-CD300lf cells (**Figure 3D and 3E**). On the other hand, disruption of the palmitoylation motif ^193^<u>CC</u>FP<u>CC</u>^198^ via mutation to ^193^<u>AA</u>FP<u>AA</u>^198^, herein Plscr1^PalmA^ mutant, significantly blunts the ability of Plscr1 to restrict MNV (**Figure 3D and 3E**)^42^. These data mirror those generated for SARS-CoV-2, suggesting that membrane association and not previously reported scramblase activity are critical for restricting MNV entry.

PLSCR1 polymorphisms have been associated with severe COVID-19 outcomes in humans^46^. Individuals with a histidine at position 262 of PLSCR1 fare better than those with a tyrosine, and this correlates with the capacity of these variants to restrict SARS-CoV-2 infection in vitro^21,22,46^. Interestingly, the equivalent position in mice, 271, is glutamine. We thus interrogated whether these polymorphisms affect PLSCR1 restriction of MNV. Human PLSCR1 containing either a histidine, tyrosine, or glutamine at position 262 all restricted MNV replication to equivalent levels (**Figure 3F and 3H**). Additionally, introduction of the human polymorphism associated with severe COVID-19 (tyrosine) into mouse Plscr1 had no detrimental impact on viral restriction **(Figure 3G and 3H)**. We were unable to assess the histidine allele in mouse Plscr1 as we could not detect stable protein. Taken together, these data suggest that Plscr1 restriction of MNV differs from that of SARS-CoV-2 as it is not sensitive to polymorphisms at the human 262 position.

## Discussion

In this study we determine that PLSCR1 inhibits the entry of MNV, a non-enveloped virus. This inhibition occurs post-attachment (**Figure 3A and 3B**) and can be overcome by a single point mutation in VP2 (**Figure 2F**). Growing evidence suggests that VP2 forms a portal that delivers viral RNA from endocytic compartments into the cytoplasm via membrane puncturing^37,38^. Thus, we propose a model in which Plscr1 may alter endocytic membranes to prevent the release of viral RNA. How VP2^Q32R^ overcomes this barrier is not clear from our data and the current understanding of VP2 oligomerization and pore formation. The Q32R substitution provides a genetic tool for investigating this dynamic yet elusive process. Overall, our findings identify PLSCR1 as a restriction factor for a non-enveloped virus and connect resistance to the VP2 membrane-penetration machinery. PLSCR1 may therefore target a host-membrane property required for both enveloped-virus fusion and non-enveloped virus genome delivery, revealing an unexpected point of convergence between distinct viral entry pathways.

Although PLSCR1 was necessary and sufficient for optimal MNV restriction in vitro, its loss did not significantly alter the overall antiviral response to type I IFN (**Figure 1H**), indicating that additional ISGs restrict MNV. Consistent with this interpretation, Plscr1-deficient mice exhibited only a modest increase in MNV^CW3^ burden in the colon, a tissue that is normally not infected by this strain (**Figure 1J**). Our cross-screen analysis also identified IFITM2, another entry inhibiting ISG, raising the possibility that PLSCR1 and IFITM2 act independently or cooperatively. Additionally, our approach to focus on genes that display antiviral activity across multiple screens deprioritized ISGs that were missed by an individual screen. Thus, *Plscr1* likely contributes to a broader IFN-induced barrier rather than acting as the major antiviral effector against MNV.

Plscr1 was initially described as a phospholipid scramblase, but this designation has been the source of debate^21,44,45^. Using previously identified loss-of-function mutations, we determine that inhibition of MNV restriction did not require a residue previously associated with scramblase activity but required an intact palmitoylation motif (**Figure 3D**). Palmitoylation of Plscr1 but not its scramblase activity, is also required for inhibition of SARS-CoV-2 by Plscr1^21,22^. In contrast to their differential effects on SARS-CoV-2, the human PLSCR1 H262 and Y262 variants restricted MNV similarly, suggesting that the requirements for PLSCR1 restriction are not identical across these viruses. However, it is possible that our assays may not be sensitive enough to detect differences in restriction by PLSCR1 polymorphisms. Previous studies of Plscr1 restriction of SARS-CoV-2 and HIV have largely focused on inhibition of membrane fusion^21-23^. However, as a non-enveloped virus, MNV entry does not proceed through membrane fusion. Our data points to a new concept in viral entry and innate immunology. Despite the diverse entry strategies of enveloped and non-enveloped viruses, host restriction factors, like Plscr1, can impose common constraints across fundamentally different classes of viruses.

## Data availability

All relevant data are within the manuscript including supporting information (**Supplementary Table 2**).

## Acknowledgements

We would like to thank Craig Wilen, Julie Pfeiffer, John Schoggins and all members of the Orchard lab for helpful discussions on this project. N.S.B. was supported by the UT Southwestern Molecular Microbiology Training Grant (T32 AI007520). R.C.O. was supported by the Burroughs Wellcome Fund Pathogenesis of Infectious Disease Program and the NIH (5R35GM142684).

## Author contributions

N.S.B. designed the project, performed experiments and helped draft the paper. L.R.P. performed experiments. R.C.O. conceptualized the project, provided supervision, and helped write the paper. All authors read and edited the manuscript.

## Materials and Methods

### Cell lines and cell culture

293T, HeLa, and BV2 cells were cultured in Dulbecco’s Modified Eagle Medium with 5% fetal bovine serum. Stable overexpression cell lines were generated by lentiviral transduction. Briefly, lentiviral plasmids were co-transfected with the packaging vector (psPax2) and the pseudotyping vector (pCMV-VSV-G) into 293T cells using PEI (Polysciences). Forty-eight hours post-transfection, lentivirus was collected, filtered through a 0.45 µm filter, and added to cells. Forty-eight hours post-transduction, media were changed to contain the appropriate antibiotic (350 µg/mL hygromycin or 5 µg/mL blasticidin and/or 1 µg/mL puromycin). All cell lines are tested regularly and verified to be free of mycoplasma contamination.

To generate HeLaΔPlscr1 cells, parental HeLa cells were cotransfected with pSpCas9(BB)-2A-Blast (a gift from Ken-Ichi Takemaru; Addgene plasmid # 118055) and pSpCas9(BB)-2A-Puro (a gift from Feng Zhang; Addgene plasmid # 62988) carrying PLSCR1 specific guides (puro: 5’ GGGTCAAGAAGTCATAACTC 3’, blast: 5’ TCAGGAGGTCTGTGATCAAT 3’). Forty-eight hours post-transfection, media were changed to contain 5 µg/mL blasticidin and 1 µg/mL puromycin for two days. Afterwards, cells were washed and returned to antibiotic free medium. Single-cell clones were isolated by limited dilution and genomic DNA was isolated (Zymo, D3025). DNA was amplified using the following forward and reverse primers (forward: 5’ ggggtatcatcaaaaactgg 3’; reverse: 5’ gcacacaacacatggaccac 3’). For each clone, indels were screened by Sanger sequencing of individual PCR reactions that were cloned in pCR-Blunt II-Topo using the Zero Blunt Topo Kit (ThermoFisher) following the manufacturer’s protocol. The resulting HeLaΔPlscr1 clone has the following alleles, indels indicated in bold with resulting stop codons in red.

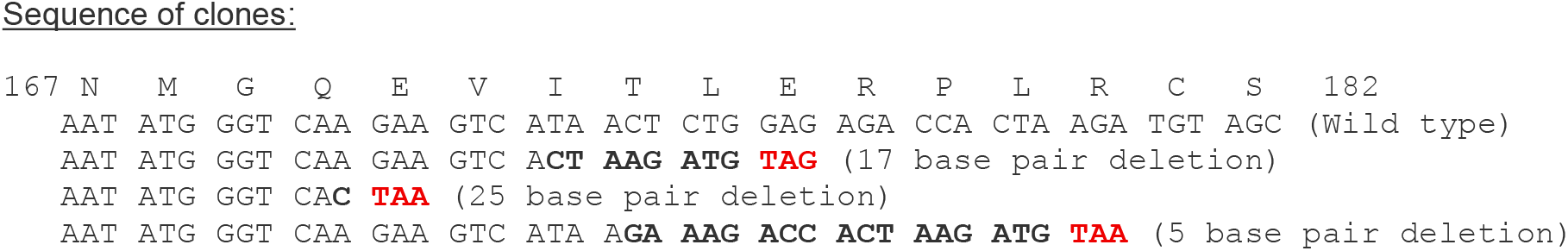

### Viral Assays

MNV^CW3^, MNV^CR6^, and respective mutants were generated by transfecting molecular clones into 293T cells and amplifying on BV2 cells as described previously^18^. For viral infections, 2 × 10^4^ indicated HeLa-CD300lf cells were seeded in 96-well plates and the next day inoculated with 25 µL MNV strains at a multiplicity of infection (MOI) of 0.05. Plates rocked for 1 hour at room temperature before adding an additional 75 µL of medium. Samples were incubated at 37°C with 5% CO_2_ and subsequently frozen at −80°C at the indicated time points. Viral titers were enumerated via plaque assay as described previously^18^. MNV-HiBiT assays were conducted as previously described^28^. Briefly, 2 ×10^4^ indicated H1 HeLa-CD300lf cells were seeded in 96-well plates and the next day inoculated with 25 µL MNV strains at a multiplicity of infection (MOI) of 0.5 or 5. Plates rocked for 1 hour at room temperature before adding an additional 75 µL of medium. Samples were incubated at 37 °C with 5% CO_2_ and subsequently frozen at −80°C at the indicated time points. After thawing, luminescent activity was measured with a Synergy LX multi-mode reader (Biotek) using a Nano-Glo HiBiT Lytic Detection System (Promega N3040) following the manufacturer’s protocols.

### Plasmid constructs

Mouse and human Plscr1 cDNA and respective mutations were cloned into pCDH-EF1-MCS-IRES-Puro lentiviral vector (System Biosciences) with an N-terminal 3xFLAG tag. Molecular clones for MNV^CW3^ (GenBank accession EF014462.1) and MNV^CR6^ (GenBank accession JQ237823) have been described previously^18^. All plasmid sequences were verified through whole plasmid sequencing prior to use.

### Immunoblotting

Samples lysed in Laemmli buffer were subjected to SDS-PAGE and subsequently transferred to PVDF membranes. Membranes were blocked in TBS-T supplemented with 5% non-fat dry milk prior to probing with antibodies. Antibodies used include mouse α-Plscr1 (PL-Scramblase 1; 1:1,000; Sigma, MABS482), mouse α-GAPDH-HRP (1:10,000; Sigma, G9295), mouse α-FLAG M2-HRP (1:2,500; Sigma, A8592), rabbit α-STAT1 phospho Y701 (1:1,000; Cell Signaling, 9167S), rabbit α-STAT1 (1:1,000; Cell Signaling, 9172S), α-rabbit-HRP (1:10,000; Thermo Fisher, 34102), and α-mouse-HRP (1:10,000; Cell Signaling, 7076S). Antibodies for MNV nonstructural proteins, including mouse α-NS1, and rabbit α-NS6/7, were used as described previously^16,47^

### Mouse infections

Animal work described in this manuscript has been approved and conducted under the oversight of the University of Texas Southwestern Institutional Animal Care and Use Committee. *Plscr1*^-/-^ mice were previously described^33^ and obtained from the European Mouse Mutant Archive (EMMA). Mice were housed in specific-pathogen-free environment at UT Southwestern. The care and use of all animals was approved by and in accordance with the UT Southwestern Institutional Animal Care and Use Committee. Plscr1^-/-^ and littermate controls were inoculated orally with 1 × 10^6^ PFU of MNV^CW3^ and immediately separated into individual cages. After 3 days of infection, mice were euthanized and tissues collected. Tissue samples were homogenized via bead beating with 1 mm silica beads (Biospec, Bartlesville, OK) in TRIzol. RNA was extracted (Zymo, R2052) and subsequently used for cDNA amplification by High-capacity-cDNA kit (Thermo Fisher, 4368813) according to the manufacturer’s protocol. Viral genome copies were assessed by TaqMan-Fast Advanced qPCR with the following probe assay: forward primer 5’ GTGCGCAACACAGAGAAACG 3’, probe 5’ [6-FAM]-CTAGTGTCTCCTTTGGAGCACCTA-[BHQ1] 3’, reverse primer 5’ CGGGCTGAGCTTCCTGC 3’. Actin copies were assessed by TaqMan-Fast Advanced qPCR with the following probe assay: Forward primer: 5’ GATTACTGCTCTGGCTCCTAG 3’, Probe: 5’ [6-FAM]-CTGGCCTCACTGTCCACCTTCC-[TAMRA] 3’, Reverse primer: 5’ GACTCATCGTACTCCTGCTTG 3’. Plscr1 copies were assessed by TaqMan-Fast Advanced qPCR with the following probe assay: Forward primer: 5’ TGCCAACTACTGATTCTTCATCT 3’, Probe5’ [6-FAM]-TGTTGTGTGTAGCTGCTGTTCCGA-[TAMRA] 3’, Reverse primer: 5’ CCATGTCTGCCCAAGTTCA 3’.

### Forward Genetic Screen for Plscr1 Resistance

These forward genetic experiments with MNV were performed under BSL2 conditions, as approved by the UT Southwestern Institutional Biosafety Committee. 2 × 10^6^ HeLa-CD300lf vector and Plscr1 expressing cells were seeded in T25 flasks and subsequently infected with MNV^CR6^ at an MOI of 0.05. Flasks were frozen after 48 hours, and subsequently clarified (10 minutes at 4,000 *g*). Fresh cells were seeded and inoculated with 1 mL of clarified virus. This passaging was repeated every 48-72 hours for eleven times. RNA was extracted from 1 mL of the final clarified virus using the Direct-zol kit (Zymo Research), and subsequently used to generate cDNA for PCR amplification. PCR products tiling the MNV genome (250 to 3000 base pairs) were sequenced using Plasmidsaurus premium PCR amplicon sequencing. Ab1 files were then aligned to the molecular clone of MNV^CR6^ WT in SnapGene, and alignment data was exported (Supplementary Table S1) to identify the percent mismatch for any given site for both vector and Plscr1 passaged virus. To account for cell culture adaptations and identify Plscr1 unique mutations, vector percent mismatch was subtracted from Plscr1 percent mismatch for each base. Mutations enriched in either population were then plotted against their respective Phred Q score, a nanopore base call confidence measurement, to account for low-quality data points.

### Entry bypass assay

4 µg of viral RNA isolated from MNV^CW3^ stocks was reverse transfected into HeLa cells stably expressing empty vector or mouse Plscr1 using lipofectamine 3000 according to manufacturer’s protocol. Importantly, these cells do not express the CD300lf receptor to prevent multicycle infections. Cells were incubated for 24 hours before plates were frozen at -80°C. Viral titers were enumerated via plaque assay as described previously^18^.

### Viral binding assay

MNV^CR6^ binding assays were performed as previously described^48^. Briefly, 2.5 × 10^5^ cells were aliquoted into cold 1.5 mL tubes on ice. 1.6 × 10^9^ PFU of MNV^CR6^ was diluted in DMEM supplemented with 5% FBS and added to each tube. Suspensions were nutated at 4°C for one hour. Cells were pelleted at 500 *g* for 5 minutes at 4°C and washed with 1 mL ice cold PBS three times before adding 300 µL TRIzol for RNA isolation and qPCR for detection of viral genome and actin copies as described above.

---

**Supplemental Figure 1:**
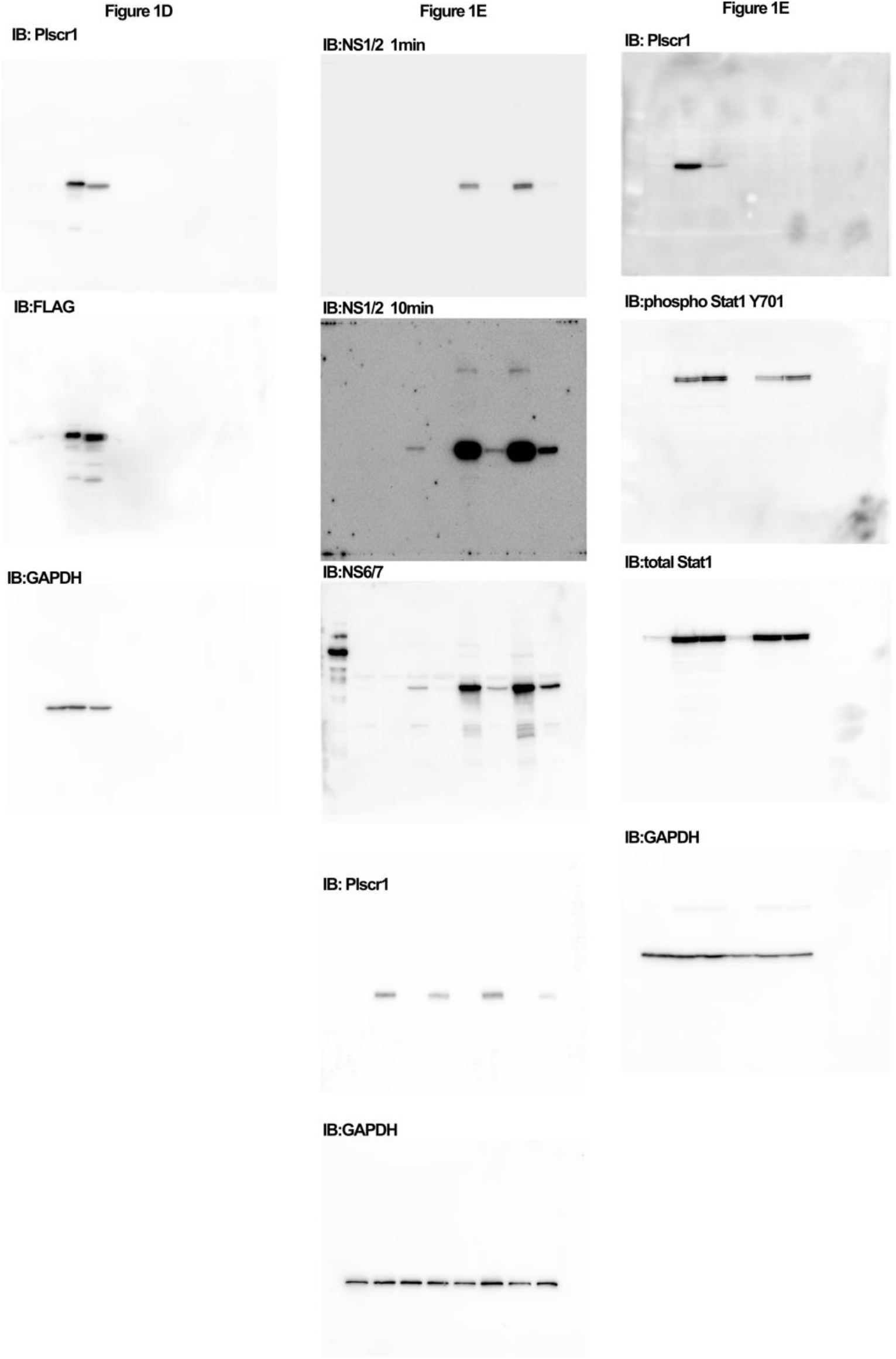
Original western Blots used in the manuscript. Unaltered immunoblot chemiluminescent images corresponding to the blots depicted in figure 1.

**Supplemental Figure 2:**
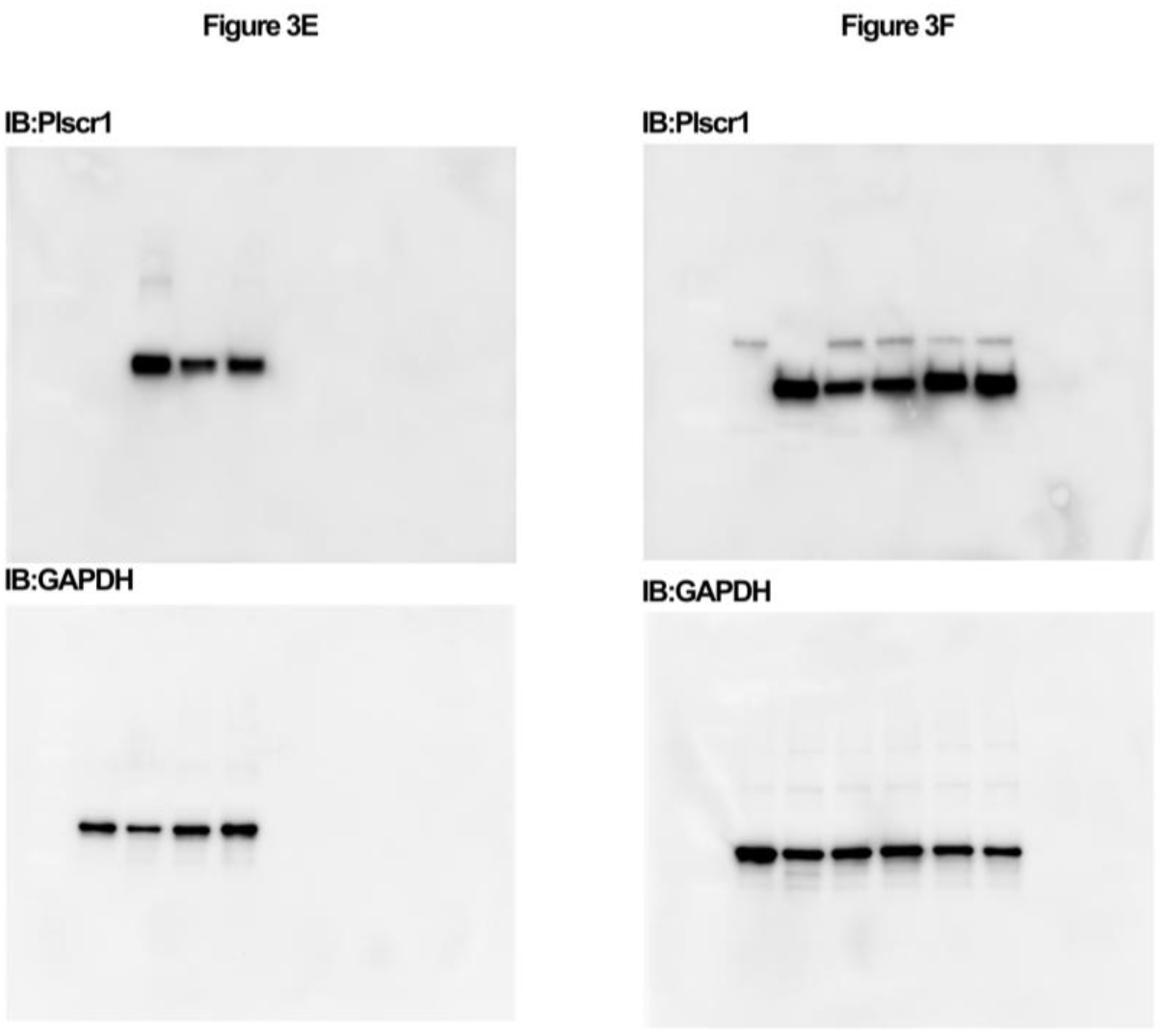
Original western Blots used in the manuscript. Unaltered immunoblot chemiluminescent images corresponding to the blots depicted in figure 3.

